# Investigating the Structural Impact and Conformational Dynamics of a Sequence Variant (c.242G>A) in *TMIE* Gene Provoking Usher Syndrome

**DOI:** 10.1101/2024.04.02.587802

**Authors:** Saqib Ishaq, Shabir Ahmad Usmani, Obaid Habib, Raheel Tahir, Abdul Aziz, Siddiq Ur Rahman, Liang Huiying

**Author notes:** contributed equally (Saqib Ishaq, Shabir Ahmad Usmani). Author contribution: First two authors.

## Abstract

Usher syndrome (USH) is a retinal autosomal recessive genetic disorder, characterized by congenital severe-to-profound sensorineural hearing loss, retinitis pigmentosa (RP), and rarely vestibular dysfunction. A transmembrane inner ear gene TMIE causing autosomal recessive usher syndrome hearing loss, which may open up interesting perspectives into the function of this protein in inner ear. This disease is linked with mutations in TMIE gene. In this study delineates the pathogenic association, miss-fold aggregation, and conformational paradigm of a missense variant (c.242G>A) resulting into (p.Arg81His) in TMIE gene segregating usher syndrome through a molecular dynamics simulations approach. The transmembrane inner ear expressed protein assumes a critical role as its helices actively engage in binding with specific target DNA base pairs. The alteration observed in the mutant protein, characterized by an outward repositioning of the proximal helical portion, which is attributed to the absence of preceding beta-hairpins in the C-terminal region. This structural modification results in the loss of hydrogen bonds, exposure of hydrophobic residues to the solvent, and a consequential transformation of helices into loops, ultimately leading to functional impairment in the TMIE protein. These notable modifications in the stability and conformation of the mutant protein were verified through essential dynamics analysis, revealing that a point mutation induces distinct overall motions and correlations between proteins, ultimately resulting in usher syndrome. The current study provides insilico evidences of Usher syndrome hearing loss disease as protein folding disorder. The energy calculation also revealed that there is a difference of −251.211Kj/mol which also indicates that the SNP has significantly decreased the stability of protein consequently folding into Usher syndrome. This study contributes molecular insights into the structural correlation between the TMIE protein and usher syndrome. The docking analysis highlight various interaction between wild and mutant structure emphasizing key residues involved in hydrogen and hydrophobic interaction.

## 1. Introduction

Usher syndrome (USH) is a genetic disorder affecting both hearing and vision. It is inherited in an autosomal recessive manner. It is considered a rare condition, that effecting approximately 1-4 individuals in every 25,000 worldwide [1] and this condition is frequently observed in Pakistani population, due to several contributing factors, including consanguineous marriages (marriages between blood relatives) and limited access to genetic testing and counseling [2]. Usher syndrome (USH) is commonly known as the “deaf–blind syndrome”, is progressive condition that usually manifest in childhood and worsens over time, leading to deafness and blindness. It is estimated that about 50% of individuals experiencing deaf-blindness are under the age of 65 [3]. British ophthalmologist Charles Usher named the disorder “Usher syndrome” in 1914, after studying a group of 69 affected individuals from 40 families, in report all showing symptoms of retinopathy and hearing loss [4]. After Pendred syndrome, USH is the most common cause of syndromic hearing loss. It is categorized into three clinical form, which is further subdivided into 14 subgroups on the basis of balance issues, the age at which visual loss first appears, and the severity of the hearing loss.

The usher syndrome is caused by mutation in any of several genes that are involved in the development and function of sensory cells in the inner ear and the retina [5-6]. It is essential to keep in mind for the purposes of this review that the condition is split into three clinical subtypes (USH1, USH2, and USH3) that were determined by clinical investigation, and that various subtypes connect to various relevant genes that were discovered by genetic investigation. To confirm or refute the clinical diagnosis, bioinformatics research is required. The identification of new genes and mutations has improved with the introduction of next-generation sequencing [7]. However, further investigations are necessary to confirm a clinical diagnosis, resulting to a large number of variations whose functions need be clarified and confirmed [8]. USH affects particular genes and proteins involved in a variety of ciliary cell processes and it can be classified as a ciliopathy. The majority of genetic mutations result in the destruction and disruption of several structural proteins that are crucial for the operation of the visual, auditory, and vestibular systems. However, there is still disagreement over whether USH qualifies as a ciliopathy.

The three main aspects of the question are terminology, histology/cytology, and pathophysiology. Due to the employment of the terms stereocilia (sing. stereocilium) and kinocilia (sing. kinocilium) to describe the structure of inner ear cells, the terminology may be a bit confusing [9]. According to histology, stereocilia composed by actin should be referred to as stereovilli (sing. stereovillum) as opposed to cilia (kinocilium) when built of tubulin (microtubules). Nevertheless, numerous pathological studies have shown that actin filaments and microtubules interact in a variety of ways, and that the appropriate functioning of actin filaments is crucial for both stereovilli and microtubules in cellular processes. The disorder’s most prevalent form, Usher syndrome, accounts for more than half of all cases [10]. It is common to describe the sensorineural hearing loss as sloping, mild to moderate in the low frequencies, and severe to profound in the high frequencies [11]. Due to the high frequency design and degree of hearing loss, infants with congenital hearing loss can be missed until the end of the first decade of life if the newborn hearing test is not accessible [12].

A variety of genetically and clinically diverse illnesses make up hereditary hearing loss. The prevalence of the most prevalent sensorineural dysfunction is approximately to be 1-3 in 1000 infants[13]. Hearing loss is caused by both X-linked dominant and recessive inheritance as well as autosomal dominant and recessive inheritance. Additionally, abnormalities that are neither monogenic or mitochondrial have also been seen in some cases. In addition to the primary phenotypes, a number of human disorders, including Usher syndrome, Alport syndrome, and Charcot-Marie-Tooth disease, frequently display hearing loss signs. At the Hereditary Hearing Loss (HHL) site, more than 120 non-syndromic hearing loss-related genes have been registered as of yet. The cause of about 75 of them is DFNB, or autosomal recessive hearing loss. Major public health concerns include hearing loss, particularly in underdeveloped nations. People with hearing loss make up two-thirds of the population in developing nations [14]. Early molecular diagnosis is crucial because prelingual hearing loss may affect how children develop language, cognition, and emotional expression.

By using whole exome sequencing (WES), this study identified the genetic origins of prelingual DFNB in five consanguineous Pakistani families. The familial genetic analysis identified four pathogenic or likely pathogenic homozygous mutations by whole exome sequencing: two splicing donor site mutations of c.787+1G>A in ESRRB (DFNB35) and c.637+1G>T in CABP2 (DFNB93) and two missense mutations of c.7814A>G (p.Asn2605Ser) in CDH23 (DFNB12) and c.242G>A (p.Arg81His) in TMIE (DFNB6). The ESRRB and TMIE mutations were novel, and the TMIE mutation was observed in two families. According to a research report, autosomal recessive non-syndromic hearing loss (DFNB) is assumed to be mostly caused by rather frequent consanguinity in Pakistan. The Whole exome sequence was identified with a disease causing mutation at (c.242G>A; p.Arg81His) in TMIE (DFNB6) family, present on the chromosome no 3. Mutation (c.242G>A; p.Arg81His) is conserved in all primates because of the conversion of Arginine into Histidine.

## 2. Materials and Methods

### 2.1. Identification of Deleterious Variant

In genetics and genomics, identifying deleterious variants is crucial and significant for understanding the potential impact of genetic variations on an individual’s health and disease risk. The identification of deleterious variants in the TMIE gene specifically aims to offer insights into potential functional effects on hearing and vestibular function. Various bioinformatics methods and tools are used for identification of deleterious variants in the TMIE gene in order to find out the impact of non-synonymous SNPs on TMIE protein. The syndrome and the TMIE gene sequence variant data was isolated from the previous research report. TMIE protein fasta sequence was retrieved from the UniProt database (http://www.uniprot.org/) accessed on 12 December 2023) through the ID: Q8NEW7. The effect of variant on the TMIE protein was recognized by following various computational tools, PROVEAN [15], PANTHER [16], SNPS&GO [17], PredictSNP [18], SNAP2 [19].

### 2.2. Secondary Structure Prediction

The reliable tool “SOPMA” server, [20] was utilized to predicted and analyzed the secondary structure elements of the TMIE gene. Sequences of the TMIE gene were obtained and submitted to the SOPMA server. SOPMA utilizes a self-optimized algorithm that combines position-specific scoring matrices, multiple sequences alignments, and statistical analysis to predict secondary structure elements. The server provided predictions for various secondary structure elements in the TMIE gene, including alpha-helices, beta-strands, and coils. To ensure accuracy, these predictions were meticulously compared with experimental data or established methods for validation. The SOPMA results were then thoroughly examined to extract the structural characteristics inherent in the TMIE gene (https://npsa-prabi.ibcp.fr/cgi-bin/NPSA/npsa_sopma.html).

### 2.3. Prediction of Gene-Gene-Interaction

GeneMANIA is a computational method used for predicting gene function and identifying potential gene interaction based on large scale of genomic data. GeneMANIA detects other genes that are associated with a conventional of input genes, using a variety of functional genomics data, including extensive database of functional association data such as protein and genetic relationships, pathways, co-expression, co-localization, and protein domains [21]. STRING (Search Tool for the Retrieval of Interacting Genes/Protein) is a biological database and online web tool that predicted protein-protein interactions, functional association and proteins based on experimental and predicted interactions. The interactions are a combination of direct (physical) and indirect (functional) connections; they are the outcome of computer prediction, information transfer across species, and interactions acquired from other (primary) databases [22].

### 2.4. Stability of Protein after Energy Minimization

The optimization 3-dimensional structure energy minimization was executed using SWISSPDB viewer [23]. Mutant and wild type structures compared for energy minimization. The detrimental effect of variant on TMIE protein stability was tested on Mupro and I-Mutant tool [24]. I-Mutant and Mupro server predict the increase and decrease stability of proteins. The altered proteins sequence were capitulated in a default parameters of I-Mutant (Temp= 25PH = 07). I-Mutant results rarely on RI (reliability index), which is usually set from 0 to 10.

### 2.5. Phylogeny Analysis of TMIE

Phylogenetic as evolutionary tree to construct among TMIE gene were compared with 30 other closed related species as identified by NCBI-BLAST program to check the maximum resemblance and confirmation of related gene family in other species. The MEGA6 program used Neighbor-Joining techniques to establish the phylogenetic relationship between proteins [25]. The interactive tree of life is an online tool for displaying, manipulating, and annotating phylogenetic and other trees (https://itol.embl.de). iTOL provides additional dataset types, more capabilities for the visualization of existing dataset types, and a number of new annotation features [26].

### 2.6. Structure prediction TMIEWT and Comparison with the TMIEMT Structure

The protein multi-template structure modeling technique was employed to model the consistent and accurate TMIEWT protein structure using the SwissModel tool [27]. Python scripts for model refinement were used to pre-refine the modeled structure [28]. The mutant 3D structure was obtained from TMIER81H protein using the amino acid swap technique in Chimera v1.15 [29]. The structure was then cleared to remove any possible clashes. Before drawing the comparison, the mutant structure was minimized with consistent configuration through GORMOS 54a7 force field [30]. Furthermore complete mutagenesis analysis and superimpose structures of wild type and mutant was performed through Biovia Discovery Studio [31].

### 2.7. Structure refinement and validation

Through Ramachandran plot [32] and ERRAT [33] 3D structure was generated and validated to check the phi and psi angles. WinCOOT v0.9.2 was used to correct the outliers, and unusual rotamers manually and remove the discontinuities and large variance in atoms B factor [34]. For more studying the validated 3D structure refine through RAMPAGE [35] and Galaxy Refine tools[36].

### 2.8. Molecular Dynamic Simulation

Molecular dynamics was run using the Desmond program for a duration of 100 nanoseconds. Hooking the protein beginning the MD modeling was the first and most crucial step in establishing the fixed viewpoint of the molecule’s binding position within the target’s active region [37-38]. MD models can simulate atom movements in a biological context and predict the binding status of receptor by mimicking their behavior over time. To optimize, minimize, and make up for any missing residues in the receptor pair, we used Maestro’s Protein Preparation Wizard. Furthermore, the System Builder tool was used in the system’s construction. The simulation was run with the TIP3P fluid model and the OPLS_2005 force field at 310 K and 1atm of pressure [39-42]. To simulate physiological conditions, 0.15 M sodium chloride and neutralizing ions were used to neutralize the models. The models were initially adjusted to feel comfortable, and every 100 ns, their progress was noted for review.

### 2.9. Molecular Docking

The TMIE protein was subjected to molecular docking using various computational tools. The ligand was selected from ChemBle, a database of bioactivity data on small drug-like compounds ChemBle [43] and was prepared using Autodock 1.5.7 [44]. The docked molecules were structurally compared using Biovia Discovery Studio [31] to investigate how two or more molecular structures fit together. This allowed us to identify the interaction of the target protein with a small molecule (ligand). The ChEMBL database is a reliable source of bioactivity data, which is regularly updated and freely available. The latest version of the database (version 22) contains comprehensive information on 1.6 million compounds, 14 million bioactivities, 11,000 biological targets, and other relevant data [43].

## 3. Results

### 3.1. Identification of Deleterious Variant

The TMIE gene sequence variants were obtained from the Ensemble Genome Browser and UniProt databases. A missense mutation (c.242G>A; p.Arg81His) was identified on chromosome 3p16.3, resulting in the conversion of arginine to histidine. To further investigate the impact of this mutation, various online tools were used for the structural and functional annotation of sequence variants. SNAP2 predicted that the mutation could affect the disease, with a score of 82 and 92% accuracy. PredictSNP also predicted a score of 87% for the mutation affecting the disease. PolyPhen2 indicated that the mutation was probably damaging, with a score of 1.000. Fathmm predicted that the mutation was damaging and could affect the protein structure, with a score of −3.24. I-Mutant predicted that the TMIE gene mutation causing deafness hearing loss would result in a large decrease in protein structure stability.

### 3.2. Secondary Structure Prediction

The SOPMA server was employed to analyzed the secondary structure compositions of both wild and mutant TMIE gene. The wild transmembrane inner ear expressed protein structure revealed a distinctive composition of 28.85% alpha helix, extended strand 25.64%, beta turn 7.69%, random coil 37.82%, other states 0.00% as shown in (Figure 1A). Conversely, the mutant structure exhibited a comparable composition with 28.21% alpha helix, extended strand 25.64%, beta turn 7.69%, random coil 38.46%, other states 0.00% as shown in (Figure 1B). Comparing the wild and mutant structure of transmembrane inner ear expressed protein, although there were minor discrepancies in the count of residues, the overall secondary structure compositions demonstrated were highly comparable. These finding indicates that the mutant within the TMIE gene has negligible impact on the overall secondary structure composition of the protein.

**Figure 1:**
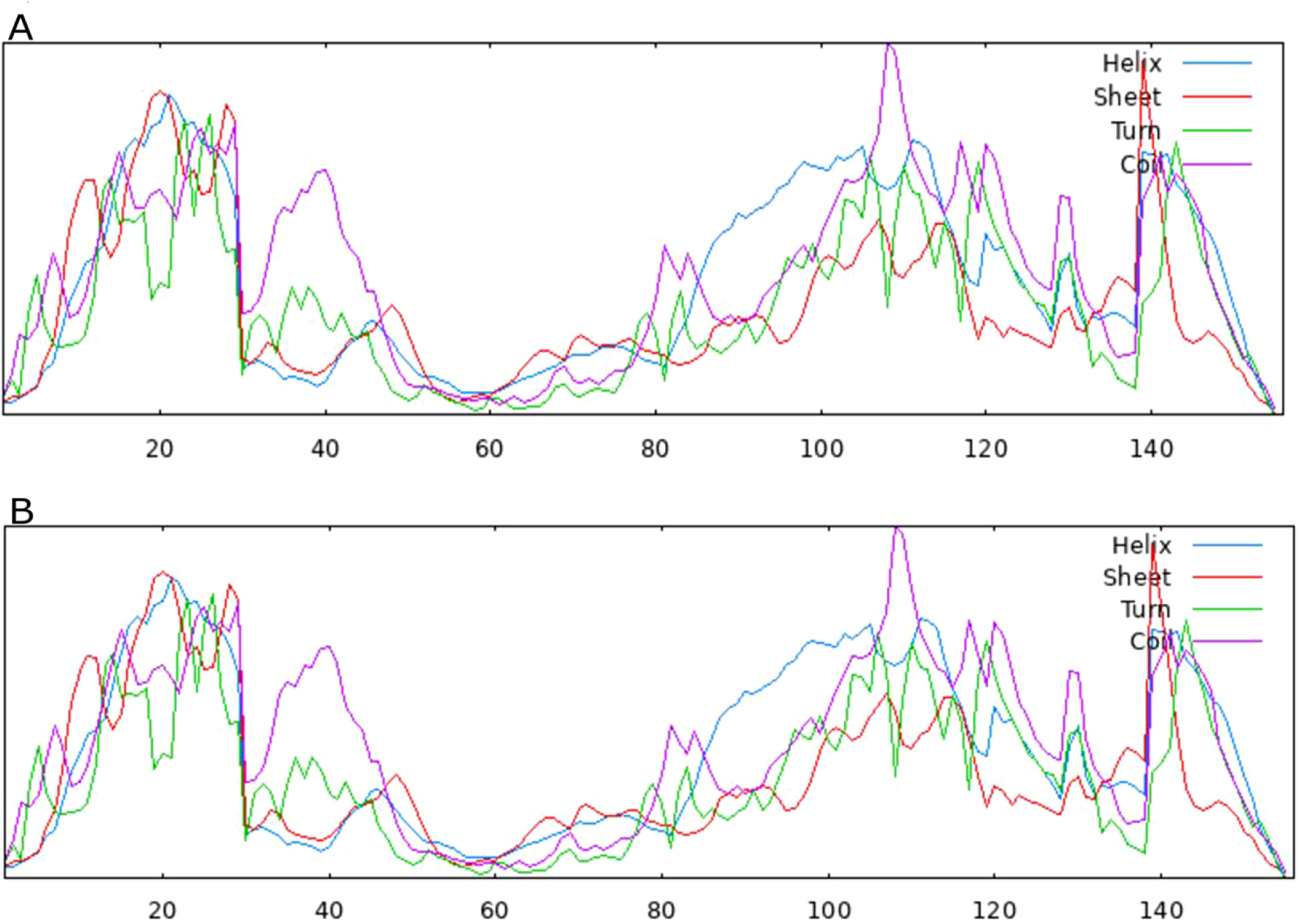
Prediction and comparison of secondary structure of *TMIE* protein.

### 3.3. Prediction of Gene-Gene-Interaction

The *TMIE* (transmembrane inner ear) networking with other genes were retrieved from GeneMANIA and STRING web servers. The GeneMANIA web source predicted functional interactions of *TMIE* (input gene) with 20 genes including, *ALS2CL, MEPE, CCL27, SIRPG, TDRD5, C5orf34, GAS8, FFAR4, WFIKKN1, MARCHF4, GPR158, C2orf81, ABCA3, CHST8, C7orf33, CNGA3, ANKRD66, C12orf65, DNAJB2*, and *CGAS* as shown in (Figure 2B). The GeneMANIA also generated the co-expression network of *TMIE* gene with all of the 20 genes. The three genes *C7orf33, TDRD5* and *DNAJB2* are found which are genetically interacted with each other. The *TMIE* gene did not show physical interaction, Co-localization, pathway, prediction, and shared protein domains networks with any of the genes. The STRING database detect the *TMIE* protein expression profile with 10 protein, *CDH23, GJB2, USH1C, STRC, LHFPL5, TMC3, STT3A, TMC2, TMC1*, and *PCDH15* (Figure 2A). The *TMIE* gene text-mining associations found with all the ten proteins while none of the active interaction (experiments, neighborhood, gene fusion, database co-occurance and co-expression) found with any of the protein. The *LHFLP5* protein is highly correlated and found with highest score among all the genes.

**Figure 2:**
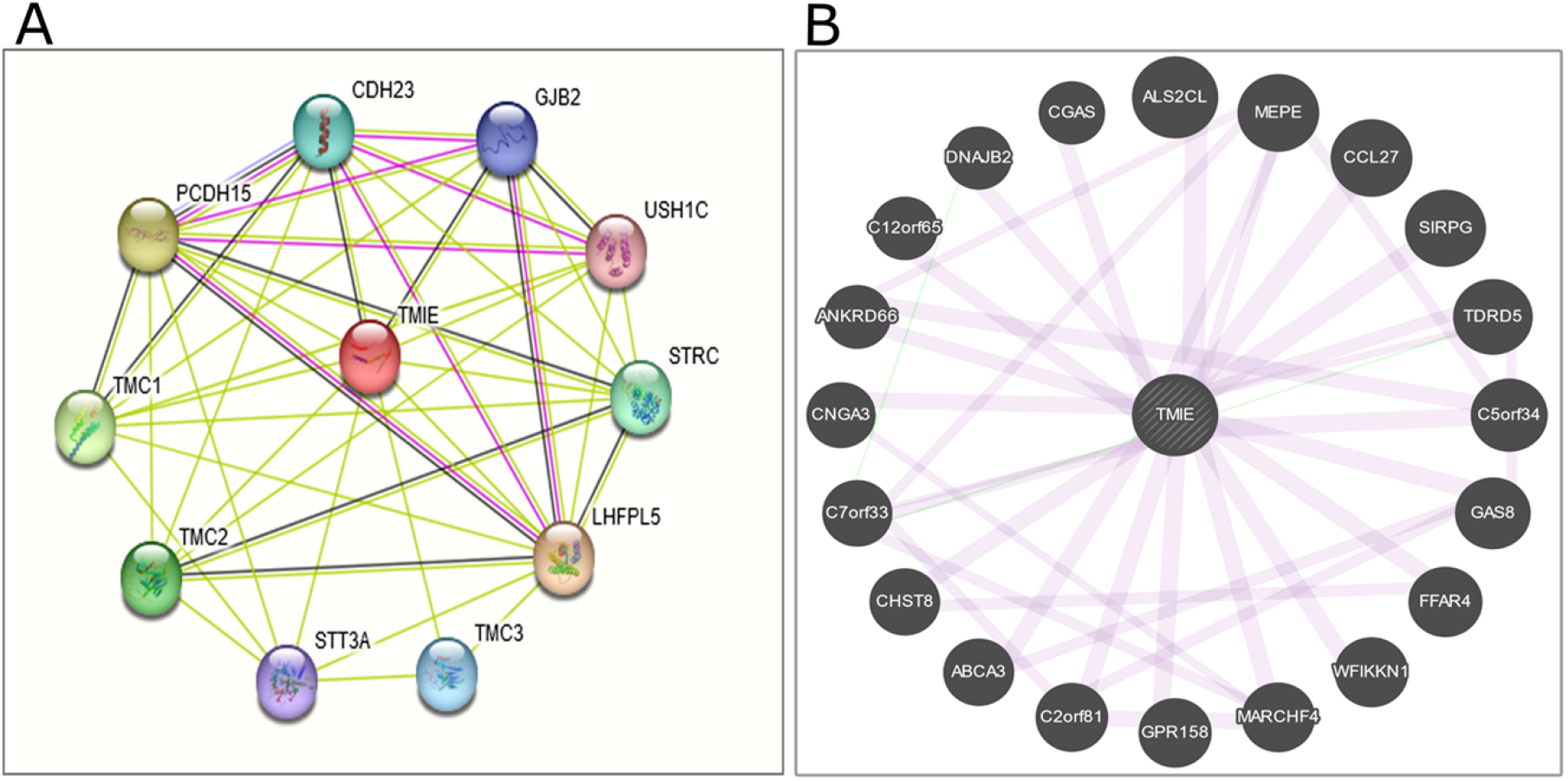
Analysis of gene-gene interaction.

### 3.4. Stability of Protein after Energy Minimization

Energy minimization was performed on both mutant and wild type *TMIE* proteins, revealing significant differences in stability between residues. The impact of amino acid alterations on protein structure stability was examined using a reliability index (RI) range of 0-10, which revealed that the *p.Arg81His* mutation was associated with decreased stability. These results propose that the mutant type is less constant than the wild type protein. (Table 1) shows the energy minimization values for bonds, angles, torsion, non-bonded, electrostatic constraints, and total values of residue to residue.

**Table 1:**
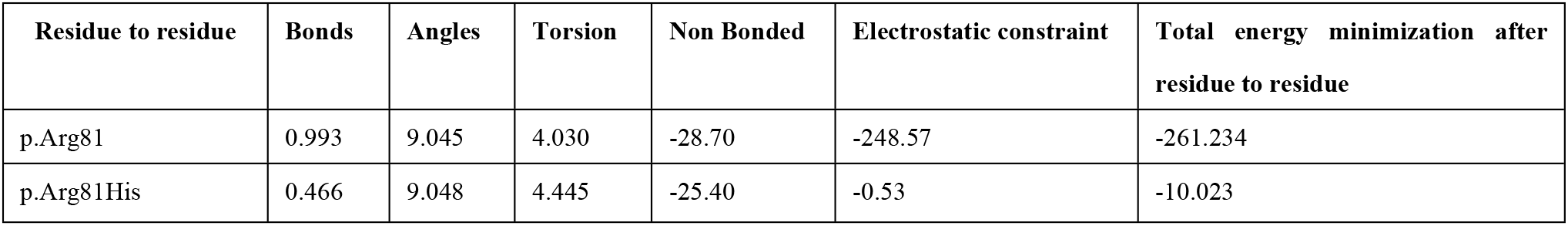
Overall energy minimization both wild and mutant *TMIE*.

| Residue to residue | Bonds | Angles | Torsion | Non Bonded | Electrostatic constraint | Total energy minimization after residue to residue |
| --- | --- | --- | --- | --- | --- | --- |
| p.Arg81 | 0.993 | 9.045 | 4.030 | -28.70 | -248.57 | -261.234 |
| p.Arg81His | 0.466 | 9.048 | 4.445 | -25.40 | -0.53 | -10.023 |

### 3.5. Phylogeny Analysis of TMIE

To confirm the conservation of *TMIE* with other species, a phylogenetic tree was constructed based on the BLAST results. MEGA6 analysis showed that human *TMIE* protein has a high degree of similarity with 30 different species, as shown in (Figure 3). The following species were found to be closely related to human *TMIE* protein: *Pan troglodytes, Papioanubis, Piliocolobustephrosceles, Macaca mulatta, Colobus angolensis palliates, Pongo abelli, Symphalangussyndactylus, Nomascusleucogenys*, and *Hylobates moloch*. iTOL provides simple manual drawing and labelling capabilities that let users add basic shapes, polygons, lines, and text labels anyplace on the tree display.

**Figure 3:**
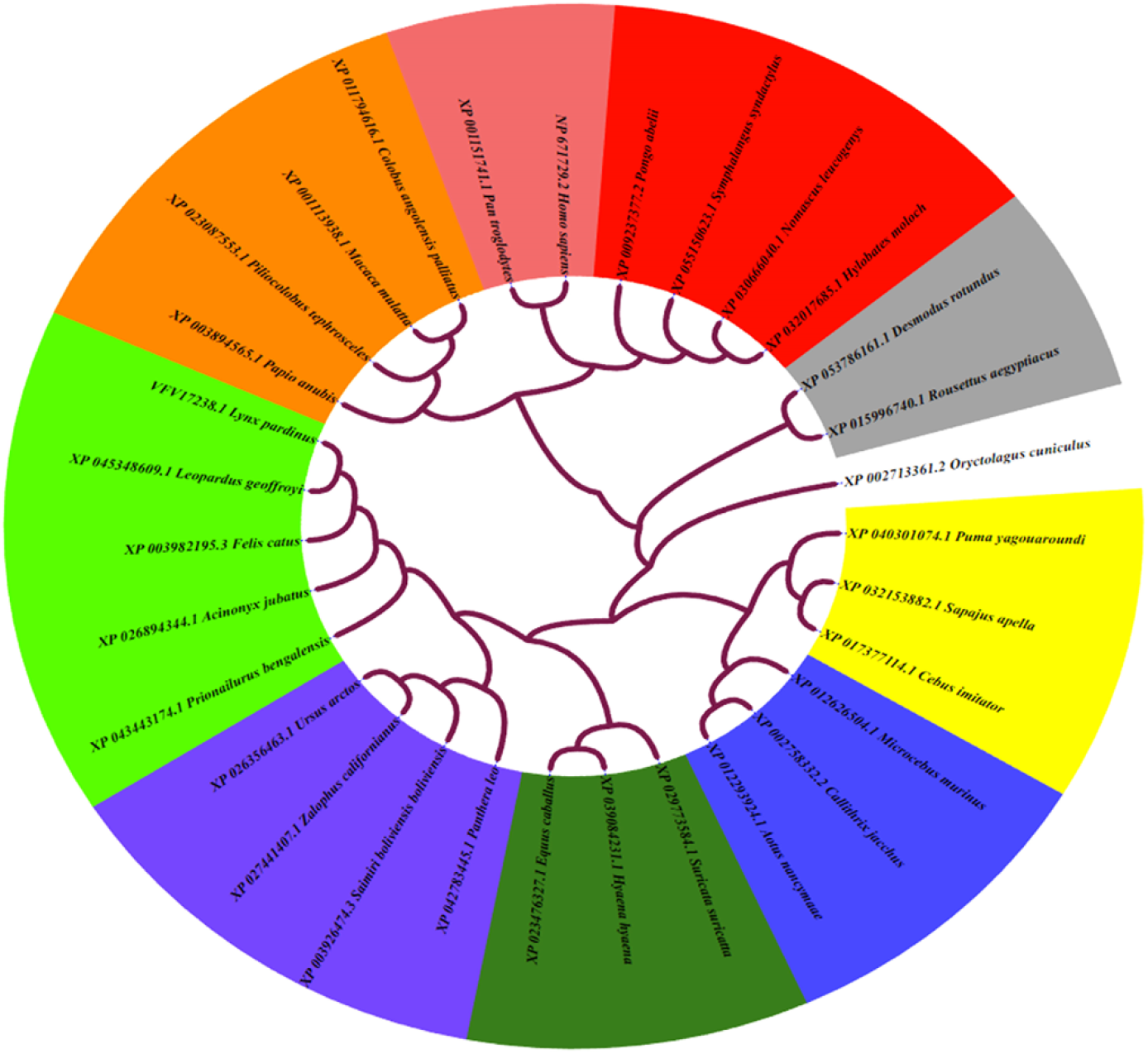
Phylogenetic analyses of *TMIE* by using the Neighbor-Joining method.

### 3.6. Structure prediction TMIEW^T^ and Comparison with the TMIE^MT^ Structure

The 3D structure of the TMIE protein was predicted using SwissModel, a web-based tool, and the resulting model was validated using various computational tools to assess different protein properties. To model the mutated structure with the p.Arg81His substitution, the “Favorite” tabs and Rotamers option of the Chimera software were used (Figure 4B). Since the predicted structure of wild TMIE protein which shows that there is maximum identity among the structure so the 3D structure we get is highly reliable, to better show the results the p.Arg81 and p.Arg81His are highlighted both wild and mutant type (Figure 4D,E). Structural comparison depicted the difference in the side chain of the residues that Arginine conserved to Histidine. TM score (template modeling) as 1.00 which shows that the structure of the two proteins share the similar folds. The TM value is among 0-1 and >0.6 means that there are 90% chances that the two protein have the same folding as shown in (Figure 4C).

**Figure 4:**
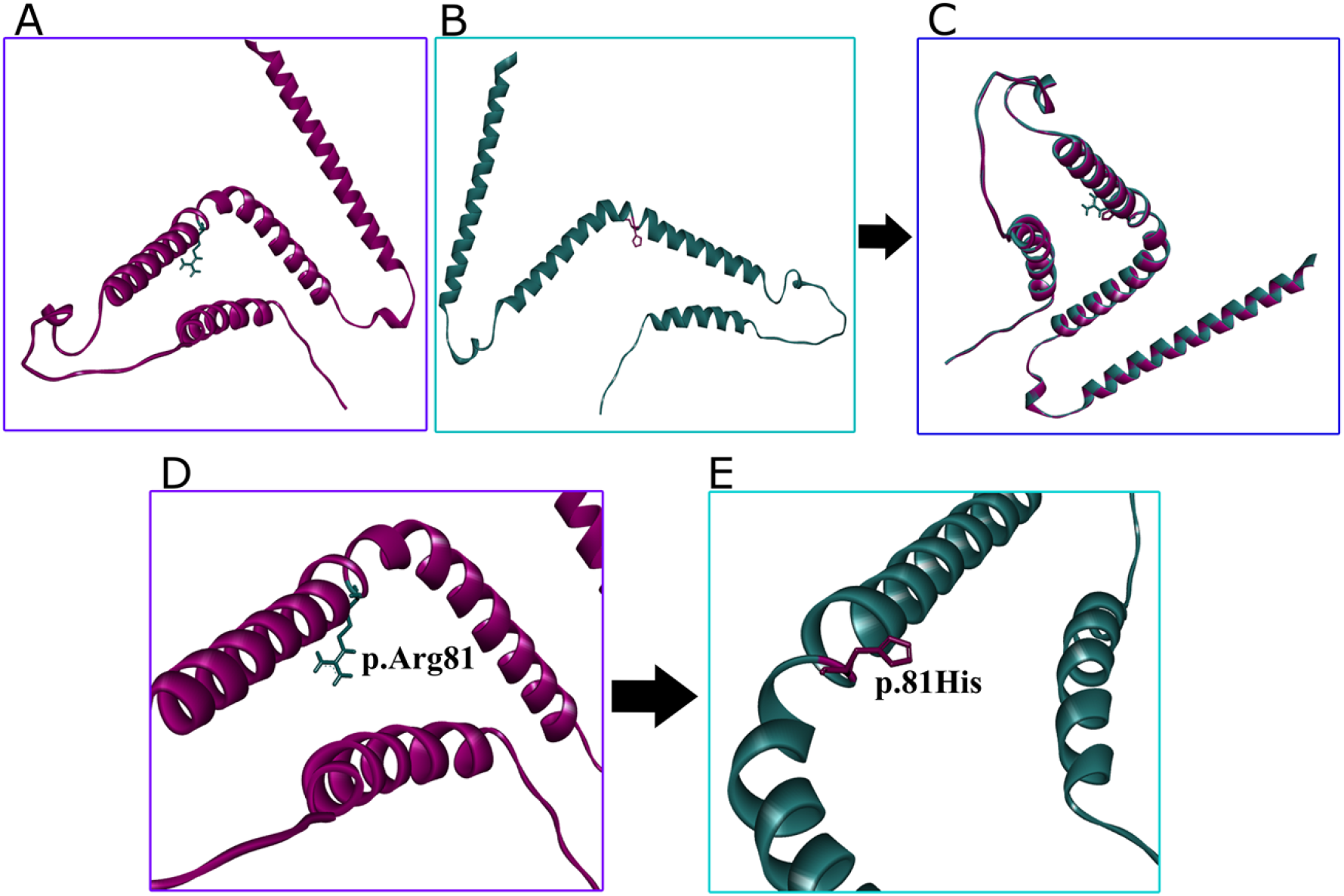
Structural analysis of wild and mutant type *TMIE* gene

### 3.7. Structure refinement and validation

The predicted 3D three dimensional structure (Figure 4A) was validated through diverse computational tools employing distinct protein properties, ensuring structure validation. The refined 3D structure was assessed using the ramachandran plot, which showed that 96.2% of residues were in the most favored regions, 3.1% were in allowed regions, and no residues were in the outlier region. The quality of the model was evaluated using ERRAT, which calculated a quality factor score of 90%. The distortion analysis of omega angles in COOT revealed the absence of unexpected peptide bond, and rotamer analysis indicated the absence of unusual rotamers in the structural conformation.

### 3.8. Molecular Dynamic Simulation

#### 3.8.1. Stability of TMIE Protein

Throughout the molecular dynamics simulation, the observed variations in RMSD values offer important information about the dynamic behavior of the wild proteins. The early RMSD variations imply that the protein was going through conformational changes and corrections as it experimented with different structural states early in the simulation. It is implied that the protein achieved a confirmation that is thermodynamically stable by the stability of RMSD values at 40 ns. Consequently, the RMSD value of TMIE^wt^ with an average of 23.99 nm through 100 ns simulation as illustrated in (Figure 5A). Analyzing the RMSD values of carbon alpha atoms in mutant proteins in figure provides intriguing insights into how these structures evolved over the course of a 100 ns simulation. These proteins stabilize at 30 ns, suggesting a rapid modification or adaption. Surprisingly, the RMSD values show a tight structural equilibrium within a 3.0 Angstrom band and are continuously stable throughout the simulation. The accuracy measured in the 3.0 Angstrom range indicates the structural stability of the mutant proteins. Furthermore, the RMSD value of TMIE^mt^ with an average of 28.89 nm through 100 ns simulation as illustrated in (Figure 5B). These findings suggest that the wild-type *TMIE* protein is more stable than the substitution R81H (Figure 5).

**Figure 5:**
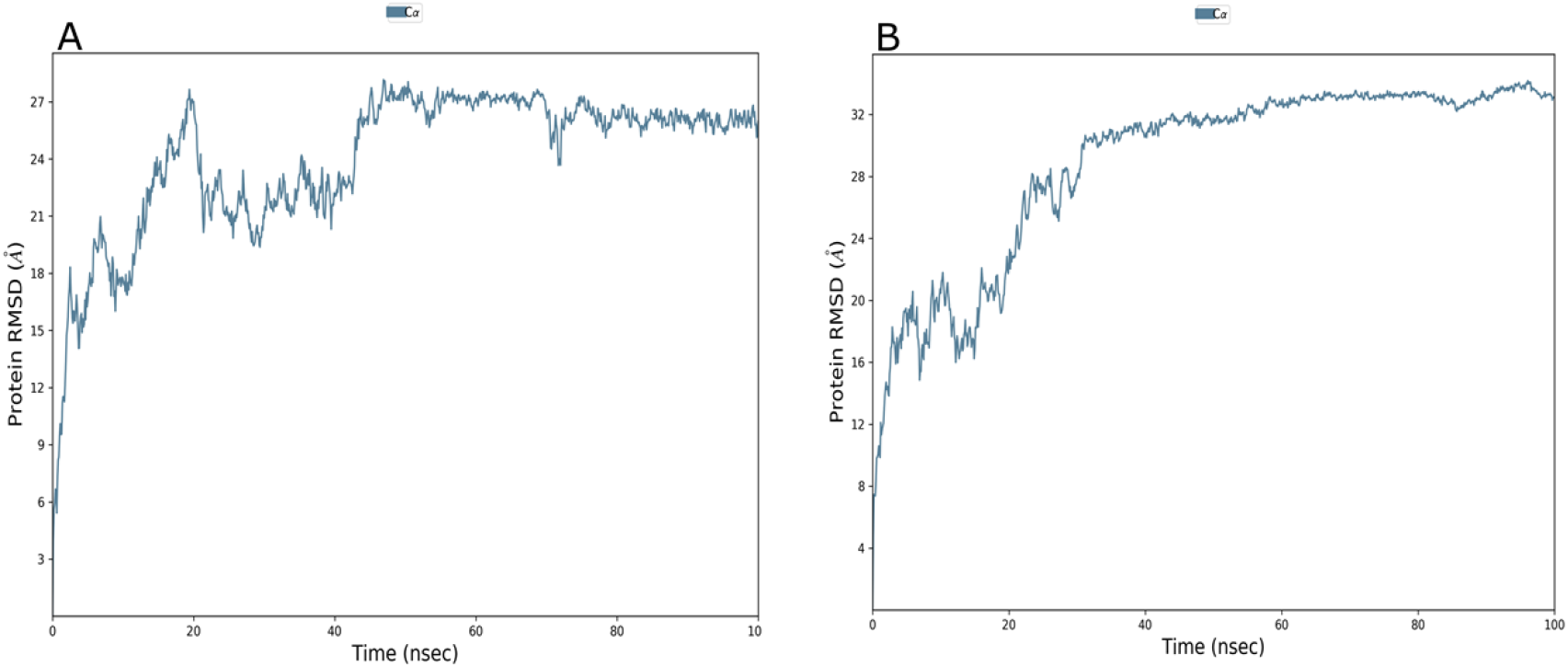
MD simulation analysis of *TMIE*^WT^ and Mutant^R81H^ proteins along the course of 100ns MD simulation. **(a)** RMSD of wt-*TMIE* **(b)** RMSD of variant R81H represents the overall instability of the *TMIE*^R81H^

#### 3.8.2. Flexibility Analysis of TMIE Protein

The residual flexibility was demonstrated by RMSF analysis as (Figure 6A,B) perceptible arginine fluctuations with substituted histidine. The variant influence the residual flexibility at 81 position along with the overall flexibility of the protein. The RMSF of wild-type arginine residue was distinguished as 10.21 nm, whereas mutated histidine was reported as 12.95 nm. The potential facilitation of this transition could be attributed to the perturbation induced by the unraveling of hearing loss with the substitution of arginine. In addition to overall rise amplitude, the RMSF in the vicinity R81H variant exhibited heightened to the *TMIE*^*wt*^.

**Figure 6:**
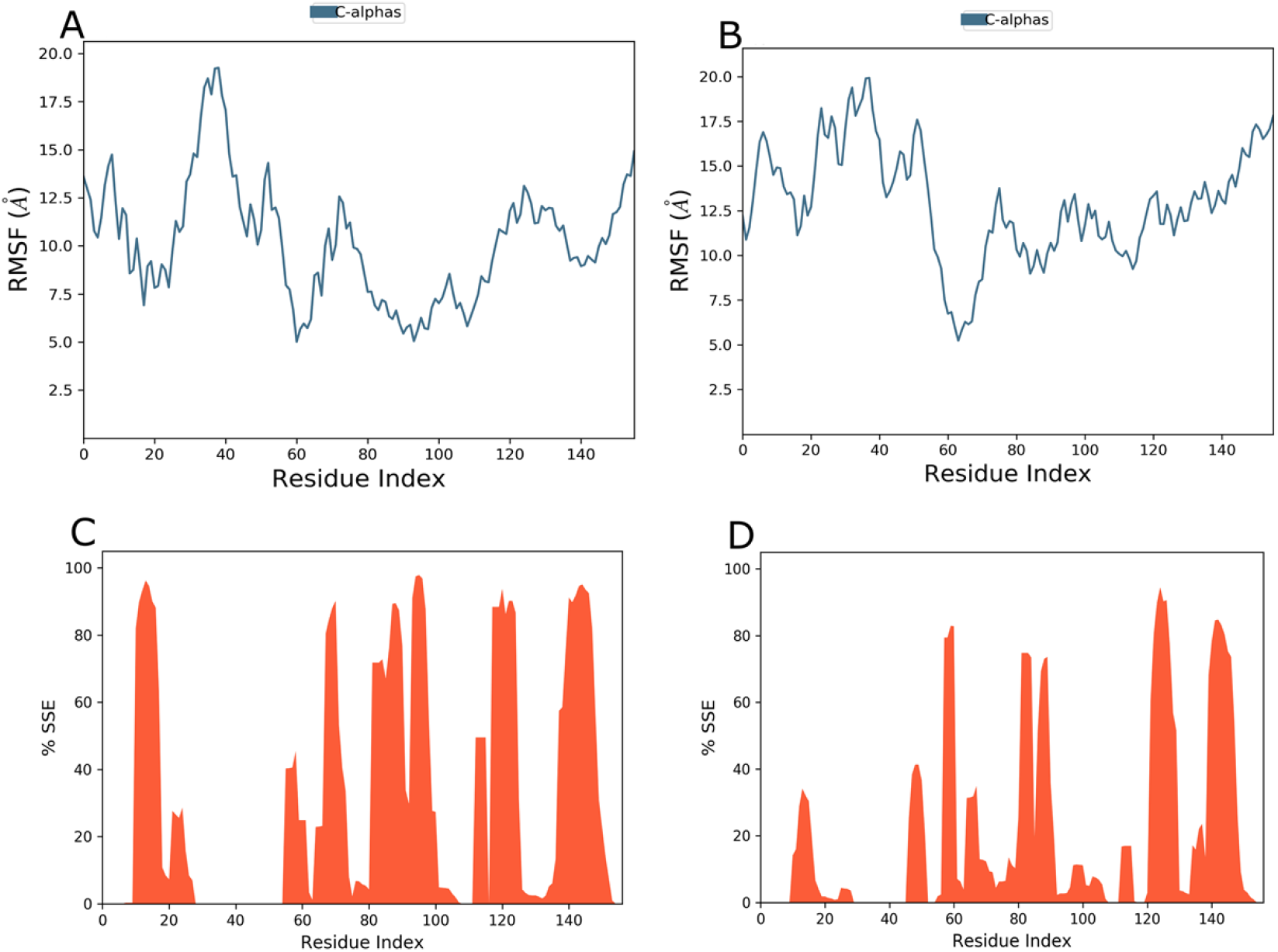
**(a)** RMSF plots for wt-*TMIE* **(b)** RMSF plots variant *TMIE*^R81H^. **(c)** The secondary structure elements (SSEs) of *TMIE*^WT^ **(d)** The secondary structure elements (SSEs) variant *TMIE*^R81H^ are also shown over the RMSF plot.

#### 3.8.3. Secondary Structural Elements

The creation of secondary structural elements (SSEs) and the structural evolution during the simulation are explained. The overall SSE percentage is significantly influenced by helices and strands. The distribution of these structural elements among residues during the simulation is shown graphically in the paper. Differential patterns in SSEs can be seen by comparing the proteins of the wild type and mutant. In the wild-type, alpha helices make up no part of the total SSE fraction, whereas beta strands make up 33.17% as shown in (Figure 6C). On the other hand, the mutant has an overall SSE percentage of 21.57%, of which 21.57% is accounted for by alpha helices and 0% by beta strands as shown in (Figure 6D). This contrast highlights important distinctions in the secondary structure composition of the mutant and wild-type proteins, offering important insights into the ways in which mutations affect the structural dynamics of the protein.

#### 3.8.4. Radius of Gyration Analysis

The gyration analysis highlights a discernible shift in the overall compactness of the R81H mutant when compared to the *TMIE*^*WT*^ structure as illustrated in (Figure 7A,B). Notably, the wild-type protein demonstrated greater compactness with an Rg score of 23.34 nm, in contrast to the mutant protein, which exhibited Rg score of 22.49 nm. This discrepancy indicates a substantial decrease in the globularity of the protein (Table 2). Over the course of the simulation, the *TMIE*^*WT*^ structure maintained compactness, while the R81H mutant displayed underscoring the mutant’s less variability. Intriguingly, the Rg value of the R81H mutant at the simulation’s onset was 18.0 nm, eventually escalating to 40.3 nm. This persistent increase suggests a loss of compactness in the mutant protein, a factor that may contribute to protein dysfunction.

**Table 2:**
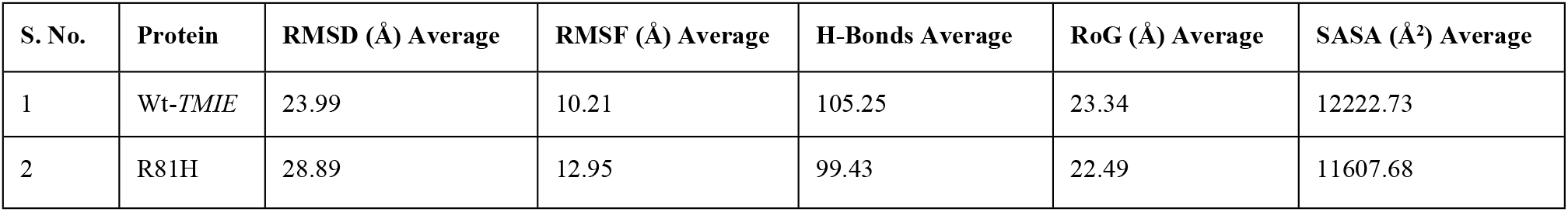
Calculated average parameters for wt-*TMIE* and mt-*TMIE* protein after 100-ns MD simulations.

**Figure 7:**
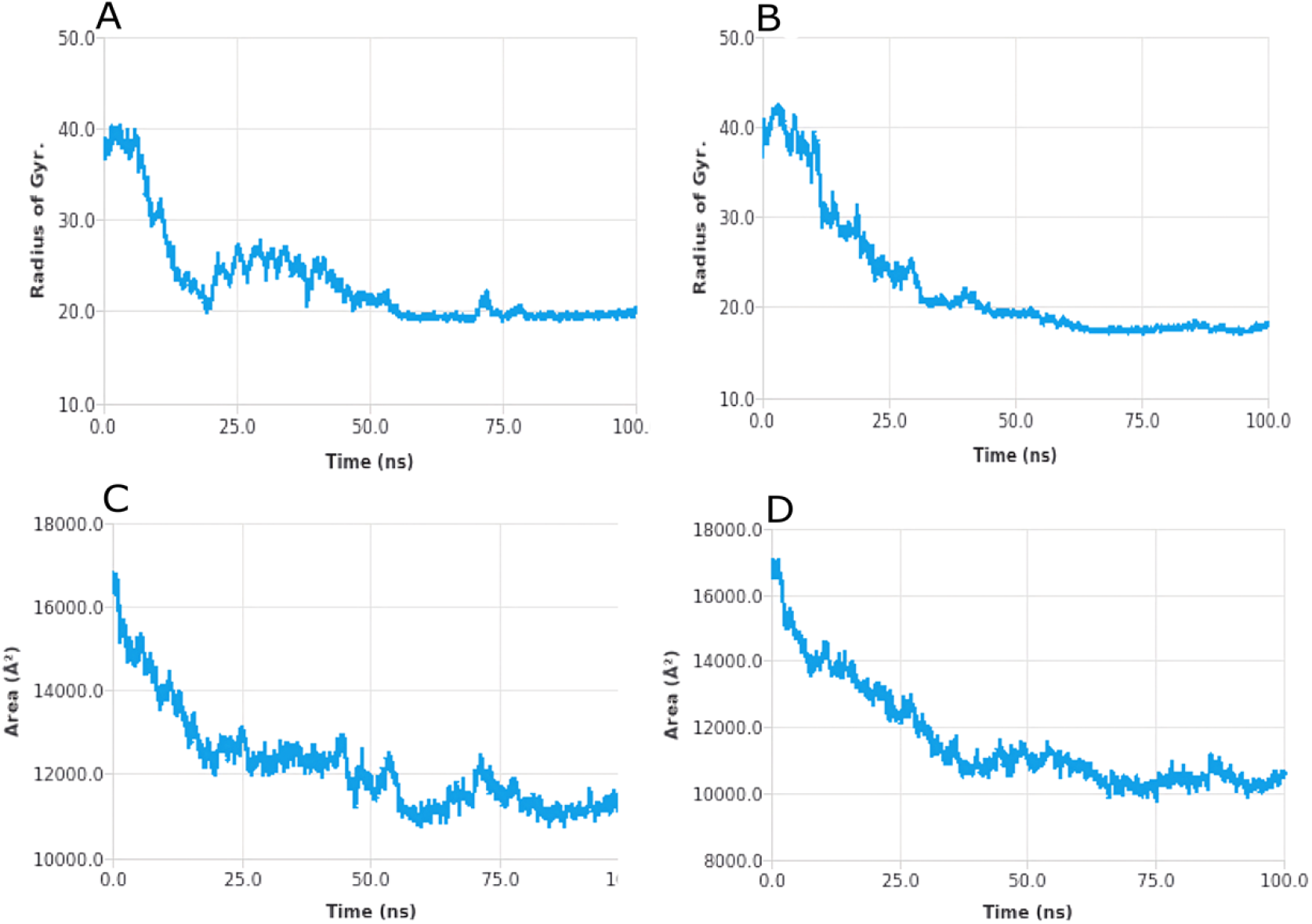
Radius of gyration, Solvent accessible surface area for *TMIE* protein shows a significant correlation between gyration and solvent accessible area. **(a)** The radius of Gyration analysis for wt-*TMIE* **(b)** And mt-*TMIE* through the course of 100 ns molecular dynamics simulation. **(c)** The solvent-accessible surface area of wt-*TMIE* **( d)** And mt-*TMIE* protein after 100 ns simulations.

#### 3.8.5. Solvent Accessible Surface Area Analysis

A comprehensive analysis of the solvent-accessible surface area (SASA) was conducted to explore the hydrophobic core regions of both the wild and mutant proteins. Notable alterations in SASA were observed for both variants, as illustrated in (Figure 7C,D). The average SASA value for the wild *TMIE* protein was 1222.73 nm^2^, whereas the mutant protein exhibited a slightly larger value of 11607.68 nm^2^ (Table 2). The substantial variance in SASA values can be attributed to the unfolding of protein folds, leading to increased exposure of the protein surface to the solvent, as depicted in figures. This exposure results in the revelation of hydrophobic residues to the solvent, ultimately causing protein unfolding and dysfunction.

#### 3.8.6. Protein Hydrogen Bond Analysis

The analysis of hydrogen bonds is essential for a comprehensive understanding of the intramolecular hydrogen bond network within the *TMIE*^*wt*^ and *TMIE*^*R81H*^ variant. (Figure 8A,B) present a detailed depiction of the H-bond network in both models, thorough examination of hydrogen bonds with a length ≤ 3.5 Å revealed significant variations in the number of hydrogen bonds with single-point mutations. Specifically, *TMIE*^*wt*^ displays an average of 105.25 H-bonds, while the *TMIE*^*R81H*^ variant exhibits an average of 99.43 (Table 2). The decrease in the number of H-bonds indicates a reduction in the stability and compactness of the protein structure. To validate the physical realism of our simulation and ensure the absence of systematic drift, we calculated and compared the total energies of both systems throughout the simulation.

**Figure 8:**
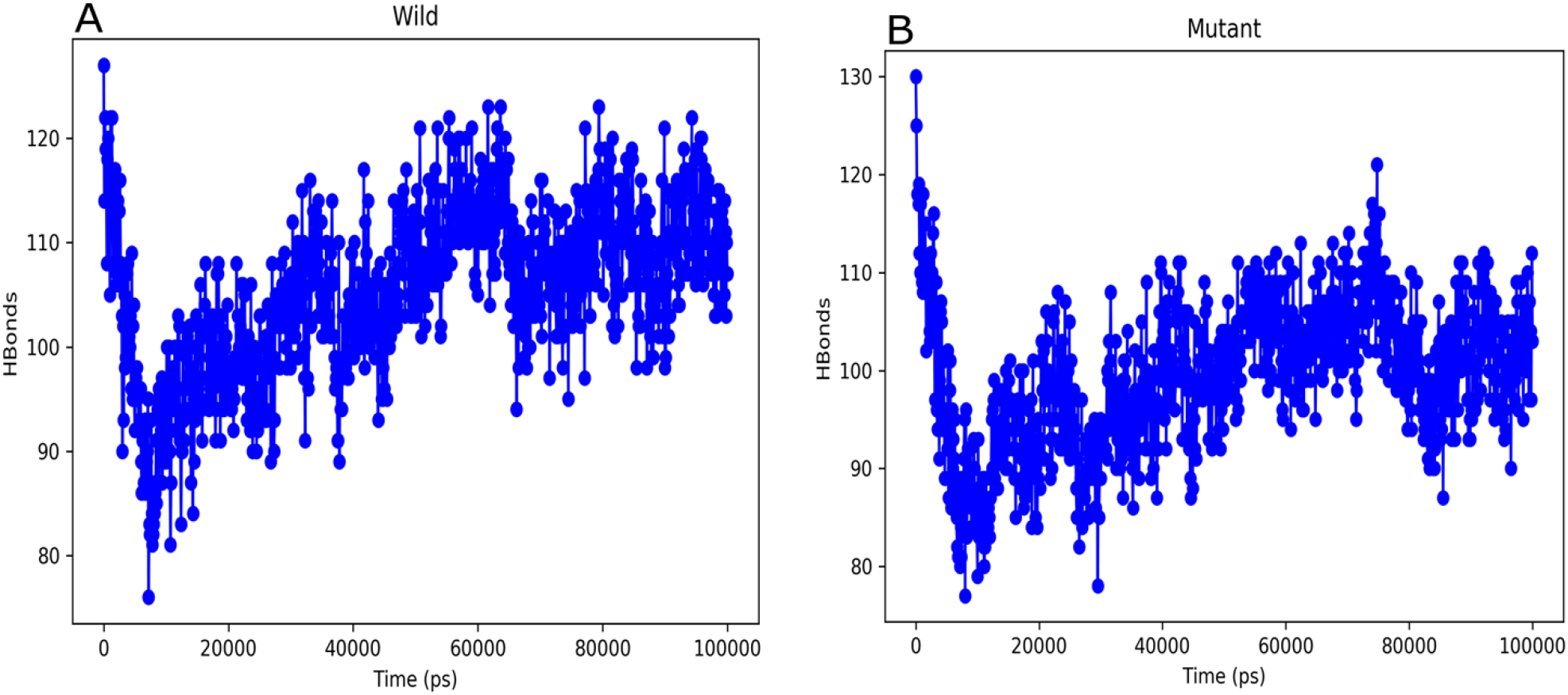
Hydrogen bond occupancy shows that the number of intra-hydrogen bonds. **( a)** The Hydrogen bond (H-Bond) of wt-*TMIE*. **(b)** And mt-*TMIE* (H-Bond) is showing an increasing trend through simulation time.

### 3.9. Molecular Docking

The acquisition of TMIE protein 3D structure was accomplished utilizing through SwissModel online web server. Subsequently, the obtained *TMIE* 3D structure underwent validation procedures and refinement processes were applied to enhance its accuracy for molecular binding interaction. Using molecular docking analysis ligand molecule retrieved from ChEMBL database (Aspartate). Selection of ligand for binding interaction with receptor on the basis of lower molecular weight (MW 414.51, MF C17H22N2O6S2). The analysis of aspartate drug ligand interacts with wild and mutant structure. Moreover, interaction of aspartate ligand of hydrogen and hydrophobic interaction with wild is Cys80, Arg81, Arg84, Thr85, Ile89, Arg92 (Figure 9A), while in case of mutant interaction with aspartate ligand is Cys80, His81, Arg84, Thr85, Ile89, Arg92 (Figure 9D). To visually represent the interaction between receptor and aspartate a create surface model was generated as shown in (Figure 9B,E). Moreover, a comprehensive examination of binding interaction between receptor and drug ligand was conducted using combination of PLIP, Pymol and Biovia discovery studio.

**Figure 9:**
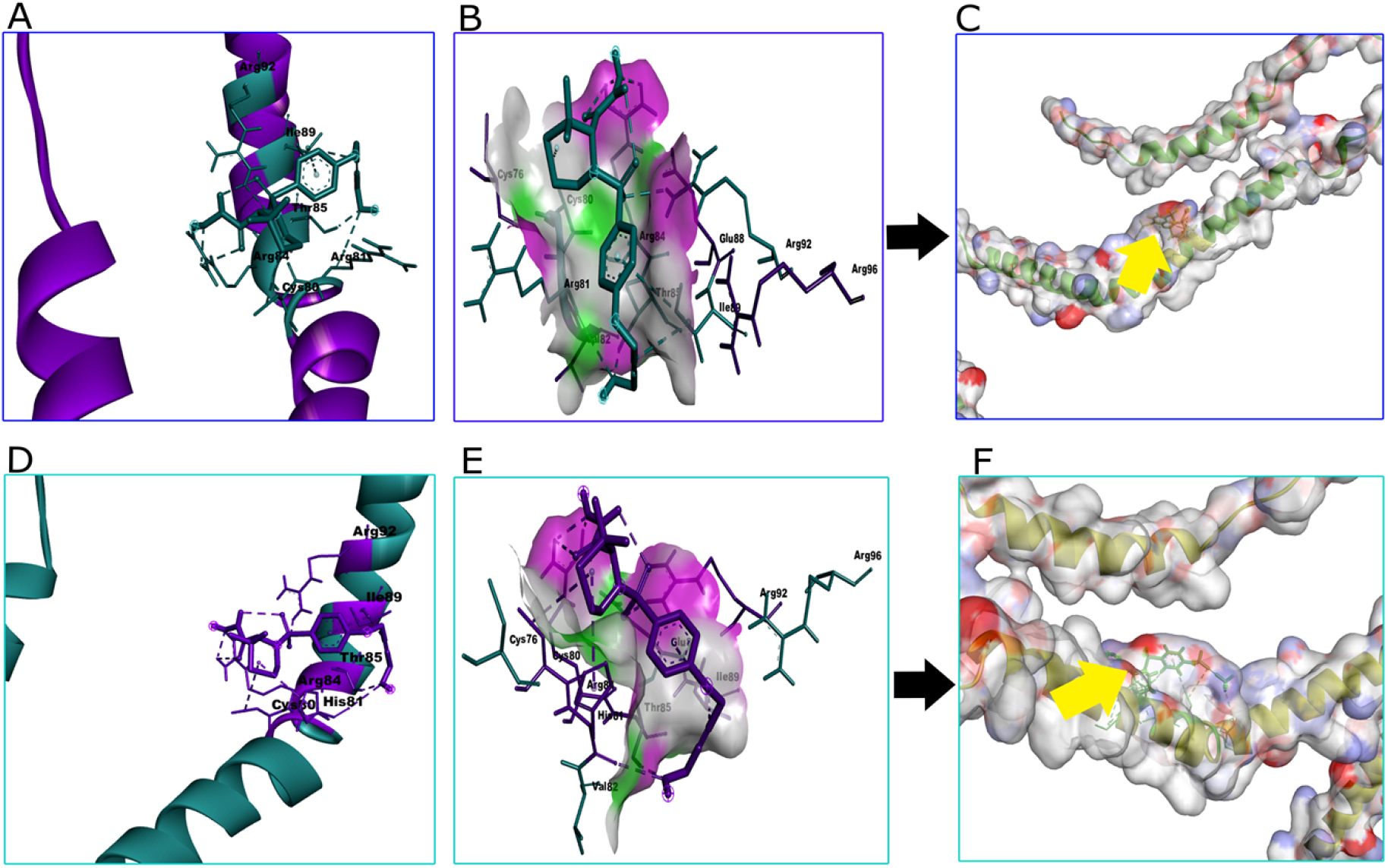
Visualization of best docked *TMIE* gene with active site binding interaction (Aspartate).

## 4. Discussion

Usher syndrome, a rare genetic disorder impacting both hearing and vision, results in deafness and blindness in affected individuals. Globally, it is estimated that Usher syndrome affects approximately 1 in 6,000 to 1 in 10,000 people [45]. The disorder is caused by mutations in several genes involved in the development and function of the sensory cells in the inner ear and retina. One of the genes implicated in Usher syndrome is *TMIE*, which encodes for a transmembrane protein that is expressed in the hair cells of the inner ear [46].

Our study is consistent with previous research that has investigated the effects of missense mutations on protein function. For example, a study on the p.R81T mutation in the *MYO15A* gene, which is associated with hearing loss, showed that this mutation led to significant structural and functional changes in the protein [47]. Another study on the p.R81G mutation in the *USH1C* gene, which is also associated with hearing loss, found that this mutation impaired the interaction of the protein with other proteins in the auditory pathway [48]. Our study provides insights into the potential functional consequences of the p.Arg81His substitution in the *TMIE* gene. Bioinformatics investigation have played pivotal role in genome wide association studies to revolutionizing the identification of mutation [49], their annotation [50], and impact [51], exploration for the discovery of novel and potent drugs [52] along with the identification of molecular targets [53].

In silico methodologies play significant role in delineating the genomic locus of gene, forecasting its transcript and elucidating its interaction with neighboring [54], discerning the structural and function attribute of protein synthesized from the gene. Utilizing in silico analysis effectively discern between deleterious and neutral single nucleotide polymorphism by leveraging diverse algorithms and information accessible in comprehensive databases [55]. In contrast to small molecules inhibitors, peptide based inhibitors exhibit reduced toxicity and enhanced selectivity for specific target. Numerous prior investigation have underscored the significance of computational method, particularly molecular dynamic simulation in expending the comprehension disease single nucleotide polymorphism across in various proteins [55].

In this study, we have established a structural association between the dysfunction of *TMIE* protein that provokes usher syndrome in hearing loss, as reported by [56]. This comprehensive study, provides a detailed exploration of the structural and molecular properties of *TMIE* to uncover its potential implications in Usher syndrome. The findings of this study encompassed meticulous structural analysis, gene mutagenesis of *TMIE*, and molecular docking analysis aimed at understanding its binding interactions with a ligand molecule. The findings also highlights the considerable impact of the p.Arg81His substitution in *TMIE* on its interactions with ligand molecules. Utilizing the SwissModel web-based tool, predicted the 3D model structure of *TMIE*, which underwent rigorous validation and refinement using various computational tools. The resulting Ramachandran plot demonstrated a remarkable structural fidelity, with 96.2% of residues situated in the most preferred regions, 3.1% in the permitted regions, and none in the outlier regions. Furthermore, the Quality factor score, determined by ERRAT, affirmed the high reliability of the predicted structure, standing at an impressive 90% and making it highly comparable to the wild-type structure. Mutagenesis analysis unveiled the alterations in the side chain of residues, particularly at position 81, indicating potential functional consequences for the protein with the substitution of histidine for arginine. Additionally, the molecular docking analysis shed light on the interactions between the aspartate ligand and both wild and mutant structures, involving crucial hydrogen and hydrophobic interactions with Cys80, Arg81, Arg84, Thr85, Ile89, and Arg92. These collective findings strongly suggest that the p.Arg81His substitution holds the potential and significance to modify the binding interactions of *TMIE* with ligand molecules, providing valuable insights into its role in Usher syndrome. The 100 ns all-atoms molecular dynamics (MD) simulation uncovered dynamic behaviors within both the wild-type and mutant *TMIE* proteins. Structural analysis revealed the wild-type *TMIE* protein’s proficient binding to the target DNA sequence through its alpha-helix. However, the introduction of the R81H mutation disrupted the protein’s overall structure, leading to the disappearance of beta-hairpins and increased flexibility. This mutations resulted in the loss of several hydrogen bonds, contributing to a reduction in compactness. A notable increase in solvent-accessible surface area ensued, exposing hydrophobic residues to the solvent and triggering protein malfunction. Essential dynamics analysis further validated these profound alterations in the stability and conformation of the mutant protein. Post-mutation, distinct overall motions and correlations between proteins emerged, ultimately culminating in usher syndrome and associated hearing loss. In addition to unraveling the structural underpinnings of *TMIE’s* association with usher syndrome, this study underscores the pivotal role of the last residue in the protein chain in correct folding, stability, and flexibility. While providing valuable insights, further studies are essential to validate and expand upon these findings. However, further studies are needed to confirm and expand upon our findings.

## 5. Conclusions

This study reports an in-silico sequence variant analysis of Usher syndrome hearing loss, revealing it to be an autosomal dominant trait. The investigation into protein folding diseases was conducted utilizing various bioinformatics tools. The sequence variant c.242G>A in *TMIE* gene resulting in the substitution of Arginine to Histidine at position 81 (p.Arg81His) has been predicted to be disease-causing by multiple bioinformatics tools. These tools use different algorithms and methods to predict the impact of amino acid substitutions on protein function and stability. Finding is consistent with the I-Mutant results, which also suggest that the variant has a destabilizing effect on the protein. It is possible that the mutation changes the location of the side chain of the affected residue from a position that is more exposed on the protein surface to a position. The structural implications of a point mutation on the *TMIE* gene were thoroughly examined through a molecular dynamics simulation assay. This study revealed the significant involvement of H1, H3, H6, and H7 α-helices in DNA binding. The consequential loss of beta hairpins resulted in structural irregularities, compactness, impacting stability, and hydrogen bonds. These deviations led to incorrect protein foldings, exposing hydrophobic residues to the solvent and ultimately causing protein dysfunction. This cascade of events contributes to the onset of usher syndrome and, consequently, hearing loss. This study elucidates the structural correlation of *TMIE*^*R81H*^ with USH, offering insight into potential contribution to the repertoire of innovative inhibitors capable competing with DNA for binding based on intrinsic features. Therefore, the overall conclusion is that this sequence variant is likely to be disease-causing in individuals with Usher syndrome.

## Author Contributions

SI, SAU, AZ, SUR: Conceptualization, Methodology, Software, Data curation, Writing-Original draft preparation. RT, OH: Helped in write-up and editing, Validation. SI, SAU: Methodology, Visualization, Validation. SUR, AZ: Reviewing and Editing, Validation.

## Acknowledgements

The authors are thankful to all members of the Department of Computer Science and Bioinformatics, Khushal Khan Khattak University, Karak, Khyber-Pakhtunkhwa, Pakistan.

## Institutional Review Board Statement

Not applicable.

## Informed Consent Statement

Not applicable.

## Data Availability Statement

Not applicable.

## Ethical approval

This article does not contain any studies with human participants performed by any of the authors.

